# GlycoTraitR: an R package for characterizing structural heterogeneity in N-linked glycoproteomics data

**DOI:** 10.64898/2025.12.16.694754

**Authors:** Bingyuan Zhang, Koichi Himori, Yusuke Matsui

## Abstract

Glycoproteomics data are rapidly accumulating due to advances in mass spectrometry instrumentation and the development of specialized search engines (e.g., pGlyco3, Glyco-Decipher) that enable identification of N-linked glycopeptide spectral matches (GPSMs) together with glycan structures. These advances have greatly expanded the scale and depth of N-linked glycopeptides; however, the intrinsic structural heterogeneity of glycosylation remains challenging to interpret. No existing tool provides a unified trait-based framework for analyzing N-linked GPSM data at both the glycosylation-site and protein levels. We developed glycoTraitR, an R package for trait-based analysis of structural heterogeneity in N-linked glycoproteomics data. GlycoTraitR provides a unified workflow to import GPSMs from search engine outputs, extract biologically interpretable glycan structural traits, and perform comparative analyses of micro- and macro-heterogeneity across experimental conditions using statistical testing.

**Implementation:** The R package and the source code of glycoTraitR are freely available on github at https://github.com/matsui-lab/glycoTraitR. A more detailed introduction and quick start guide are avaible at https://matsui-lab.github.io/glycoTraitR/.

## Introduction

Recent advances in both mass-spectrometry instrumentation and computational methods have substantially expanded the depth and reliability of glycoproteomics (Khoo, 2021). Modern search engines such as pGlyco3 and Glyco-Decipher further increase confident N-linked glycopeptide spectral matches (GPSM) through “glycan-first” strategies with rigorous false-discovery-rate (FDR) control. However, GPSM interpretation remains challenging: current analyses typically focus on individual glycopeptides and do not fully leverage the structural information encoded in the attached glycans. As a result, the field lacks a reproducible, systematic framework for analyzing N-linked glycoproteomics data.

We focus on a fundamental yet often under-analyzed feature of glycoproteins: their intrinsic heterogeneity. Glycoprotein heterogeneity occurs at two levels (Potel et al., 2025). Macroheterogeneity refers to the diversity of glycan structures attached across the glycosylation sites of a protein, whereas microheterogeneity captures structural variation among glycoforms at a single site. These heterogeneous glycoforms influence protein folding, stability, receptor binding, and downstream signaling. Large-scale glycoproteomics studies have demonstrated that such site-specific heterogeneity is extensive and reproducible, and that glycan composition patterns and site-specific glycan ratio changes can sensitively capture biologically relevant variation (Potel et al., 2025; Wu et al., 2025; Zhu et al., 2025).

However, there is no analytical framework for quantifying and testing macro/micro-heterogeneity change in N-linked glycoproteomics data. Recent trait-based approaches developed in the glycomics field--such as glycowork (Thomès et al., 2021), glycompare (Bao et al., 2021), and glytrait (Fu et al., 2025)—demonstrate that analyzing glycan substructures can reveal meaningful biological patterns. However, these methods focus on glycan and do not incorporate peptides sequence context. Consequently, no tool provides a unified trait-based framework for N-linked GPSM data that integrates glycan structure with peptide sequence. To address this gap, we developed an end-to-end analytical framework for quantifying macro- and microheterogeneity directly from N-linked GPSMs, enabling interpretable trait-level comparisons at both the site and protein levels. We implemented this framework in the R package glycoTraitR, which integrates with pGlyco3 and Glyco-Decipher outputs and supports trait extraction, differential analysis, visualization, and user-defined motifs for flexible, biologically driven analyses.

## GlycoTraitR Pipeline Overview

GlycoTraitR implements a workflow that transforms N-linked glycopeptide spectral matches (GPSMs) into quantitative measures of glycan structural heterogeneity at both the glycosylation-site and protein levels. As summarized in Figure 1, the pipeline takes GPSM files produced by the glycoproteomics search engines, derives structural traits for each N-glycan, constructs a trait--GPSM count matrix, and then aggregates traits to site and protein levels for statistical testing between experimental conditions.

**Fig. 1.**
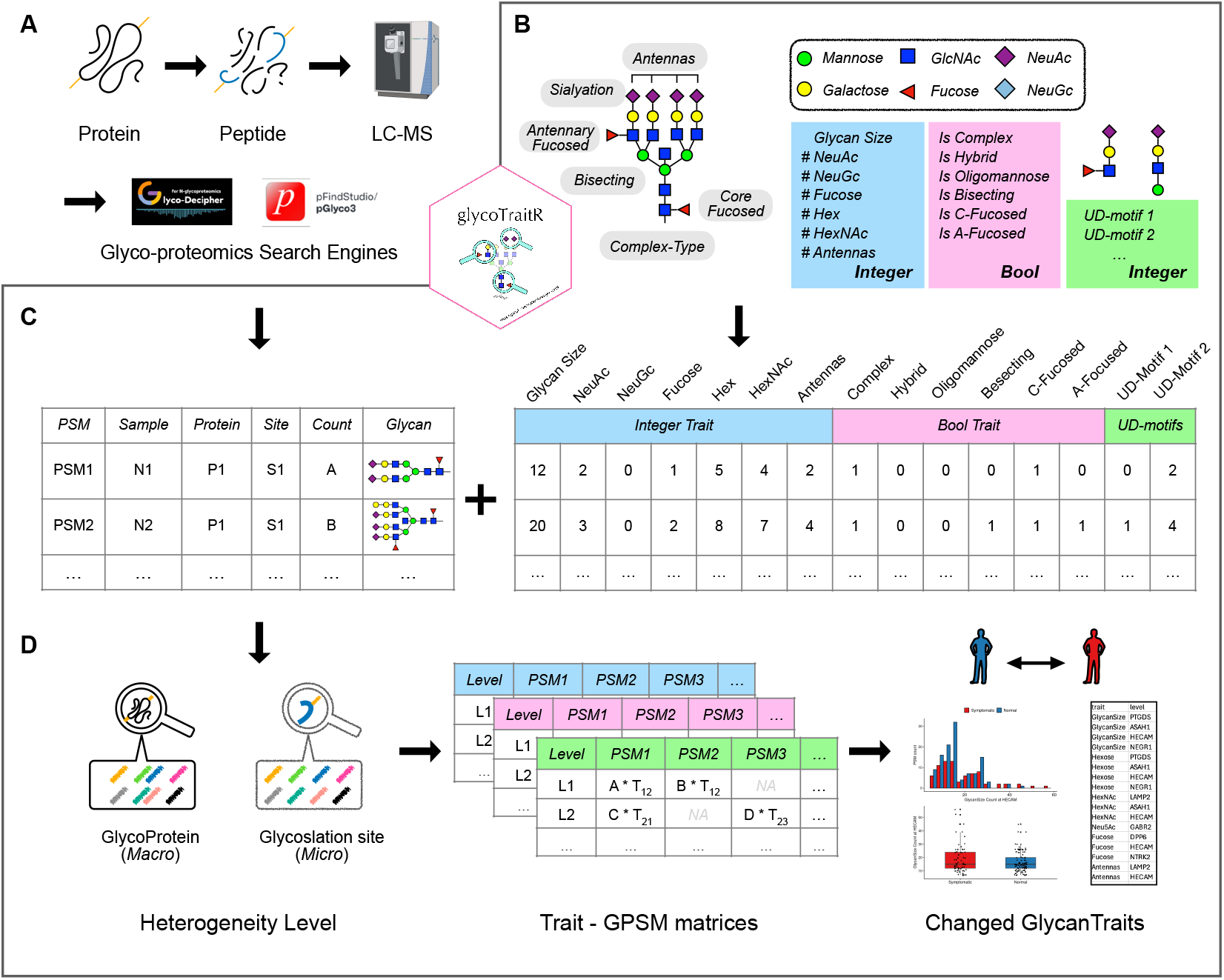
Overview of the glycoTraitR workflow. (A) LC--MS/MS glycoproteomics experiments generate peptide spectra that are searched with glycoproteomics engines such as pGlyco3 and Glyco-Decipher, yielding N-linked glycopeptide spectral matches (GPSMs) with assigned glycan structures. (B) For each distinct N-glycan, GlycoTraitR parses the structure and decomposes it into a panel of compositional and structural traits. Moreover, user-defined glycan motif (UD-motif) can be specified as additional traits. (C) The traits are joined back to the GPSM table to construct a trait-annotated GPSM matrix, containing integer-valued traits and Boolean traits for each GPSM. (D) GlycoTraitR aggregates the trait--GPSM matrix to the glycoprotein level (macroheterogeneity) and glycosylation-site level (microheterogeneity), performs trait-wise comparisons between groups. The resulting trait-level statistics and visualizations highlight traits whose abundance or dispersion changes.

### Input

GlycoTraitR accepts a table of N-linked GPSMs generated from LC–MS/MS data and processed by glycoproteomics search engines that report glycan structures, such as pGlyco3 or Glyco-Decipher. The table contains a unique GPSM identifier, sample label, protein accession, glycosylation-site, glycan-structure string, and a numeric spectral-count column used to quantify each GPSM (Figure 1A). In addition, metadata linking sample labels to group labels is required.

### Trait Definition

For each assigned N-glycan, GlycoTraitR parses the reported glycan string and decomposes it into a predefined set of structural traits that capture biologically interpretable aspects of glycan composition and architecture. The parser directly reuses glycan annotations produced by glycoproteomics search engines, enabling consistent trait extraction without redefining glycan structures. Both composition-based and structure-based traits are supported, allowing glycan heterogeneity to be represented at the trait level rather than as individual glycoforms (Figure 1B).

### User-defined traits

In addition to the built-in trait set, GlycoTraitR allows users to define custom glycan motifs as user-defined traits. These motifs are evaluated for each GPSM and incorporated into subsequent analyses, enabling flexible, hypothesis-driven interrogation of specific glycan substructures.

### Trait summarization at protein and site levels

Trait values are merged with the GPSM table to generate a trait-annotated representation of each GPSM (Figure 1C). Depending on the analysis objective, traits are aggregated either at the protein level, capturing macroheterogeneity across glycosylation sites, or at the site level, capturing microheterogeneity among glycoforms at individual sites, and used for downstream comparison across experimental conditions.

### Statistical comparison and output

Aggregated trait values are compared across experimental groups using standard statistical tests to assess differences in trait abundance and variability at both the site and protein levels. Results are summarized in trait-level tables and visualizations, enabling direct interpretation of changes in glycan micro- and macroheterogeneity (Figure 1D).

## Discussion

In this work, we present a trait-based framework for quantifying glycan structural heterogeneity in N-linked glycoproteomics data. By decomposing assigned N-glycans into biologically interpretable structural traits and aggregating these traits at the glycosylation-site and protein levels, glycoTraitR enables systematic evaluation of micro- and macroheterogeneity across experimental conditions. This representation shifts the analytical focus from individual glycopeptide-level comparisons to trait-level patterns, facilitating scalable and interpretable analyses of complex glycoproteomics datasets.

The current implementation of glycoTraitR relies on discrete N-glycan structures reported by glycoproteomics search engines such as pGlyco3 and Glyco-Decipher and uses GPSM counts as quantitative measures. Consequently, the framework inherits the inherent trade-offs of tandem mass spectrometry–based structure assignment, where identification depth, structural resolution, and confidence must be balanced. As additional search engines begin to report explicit and standardized glycan structures, trait-based frameworks such as glycoTraitR will support more systematic cross-study comparisons and integrative analyses. Currently, glycoTraitR takes GPSM count value as the input. As a future work, it can be extended to take intensity-based quantification output as the input once the quantification output with glycan structures becomes available for N-glycoproteomics search engines.

The predefined trait library in glycoTraitR focuses on monosaccharide composition and canonical architectural features, while support for user-defined motifs allows flexible extension toward study-specific or functionally motivated substructures. As functional annotations and biosynthetic knowledge of N-glycans continue to expand, the trait space can be further enriched with topology-aware or data-driven features. In this context, glycoTraitR provides a practical and extensible foundation for trait-centered interrogation of N-glycosylation heterogeneity.

## Supporting information

Supplementary_glycoTraitR

