## Supplementary_glycoTraitR for "GlycoTraitR: an R package for characterizing structural heterogeneity in N-linked glycoproteomics data"

### Abstract

In this supplementary, we introduce the definition of default trait and include a case study of application for glycoTraitR package to Alzheimer’s disease brain glycoproteomics data. These results demonstrate that trait-based glycan representation enables robust and biologically meaningful comparison of N-glycosylation patterns.

### Default Trait Definition

GlycoTraitR includes a set of predefined traits that capture a glycan's compositional and structural features. These traits are derived by parsing the monosaccharide composition and structure of the glycan string from either WURCS 2.0 or pGlyco3 format. The default trait library is designed to convert different glycoforms into common biologically interpretable descriptors, enabling quantification of N-glycan heterogeneity.

The default traits fall into two categories: (i) integer traits and (ii) Boolean traits. This representation follows established and known characterizations of N-glycans, including glycan class (oligomannose, hybrid, complex), branching degree, core fucosylation, antenna fucosylation, and bisecting N-acetylglucosamine. Together, these integer and Boolean traits provide a compact yet expressive representation of each N-glycan for downstream quantitative analyses. The default traits are defined as follows:

*Integer traits*

- Glycan size: Total number of monosaccharide residues.
- HexNAc: Number of N-acetylhexosamine residues.
- Hex: Number of hexose residues (mannose or galactose).
- Fucose: Number of fucose residues.
- NeuAc: Number of N-acetylneuraminic acid residues.
- NeuGc: Number of N-glycolylneuraminic acid residues.
- Antennas: Represents the number of terminal branches extending from the Man3GlcNAc2 core. We define an antenna as a branch that originates from one of the mannose residues extending outward from the core and is initiated by an N-acetylglucosamine (GlcNAc) residue.

*Boolean traits*

- IsComplex: A glycan is defined as a complex-type N-glycan when having two or more branches extending from the central mannose residues Man1GlcNAc2, and these branches are elaborated with N-acetylglucosamine units rather than mannose alone.
- IsHybrid: A glycan is defined as a hybrid N-glycan when containing features of both oligomannose and complex-type structures: at least one branch retains a mannose-only extension, while another branch is extended with N-acetylglucosamine or other complex-type residues.
- IsOligomannose: A glycan is considered oligomannose when it contains multiple mannose residues extending beyond the conserved Man3GlcNAc2 core, and all outward branches arising from the central mannose unit consist exclusively of mannose.
- IsBisecting: A glycan with the a bisecting N-acetylglucosamine (GlcNAc) residue attached to the central mannose (Man1GlcNAc2).
- IsC-Fucosed: A glycan with a core fucosylation.
- IsA-Fucosed: A glycan with at least an antenna fucosylation.

These traits provide a compact yet expressive representation of N-glycan that is suitable for quantitative comparisons. User-defined motifs are treated as integer-valued traits, and their values corresponding to the number of times each motif occurs within the glycan structure.

*Example:*

We provide an example of how glycoTraitR parses a N-glycan into traits. The glycan (G42295MA) from glycosmos (Yamada et al., 2020) is shown in Figure S 1.

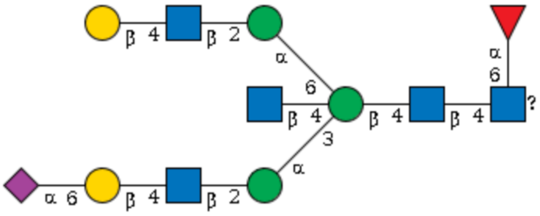


Figure S 1. The example N-glycan (G42295MA)

The following is the wurcs 2.0 format of the glycan, and we demonstrate how to extract its structural traits using glycoTraitR functions.

```
# WURCS representation of the glycan
wurcs <- "WURCS=2.0/7,12,11/
 [a2122h-1x_1-5_2*NCC/3=O][a2122h-1b_1-5_2*NCC/3=O]
 [a1122h-1b_1-5][a1122h-1a_1-5][a2112h-1b_1-5]
 [Aad21122h-2a_2-6_5*NCC/3=O][a1221m-1a_1-5]
 /1-2-3-4-2-5-6-2-4-2-5-7
 /a4-b1_a6-l1_b4-c1_c3-d1_c4-h1_c6-i1_d2-e1_e4-f1_f6-g2_i2-j1_j4-
k1"
```

```
glycanglycan_tree <- glycoTraitR:::wurcs_to_tree(wurcs)
glycoTraitR::compute_glycan_traits(glycan_tree, motifs = NULL)
```

```
GlycanSize    Hexose      HexNAc     Neu5Ac      Neu5Gc      Fucose
     12          5           5          1           0           1
Antennas      Bisect     Complex     HighMan      Hybrid     CoreFuc
     2           1           1          0           0           1
AntFuc
     0
```

### Case Study

Next, we showcase and apply glycoTraitR to a publicly available dataset. The case study follows the complete workflow in glycoTraitR, including data import, trait derivation, trait–GPSM matrix construction, statistical testing, and visualization.

### Data

The case study is based on the intact N-glycoproteomics dataset (Suttapitugsakul et al., 2022). The full dataset is publicly available under the ProteomeXchange identifier *PXD032219*. It consists of LC-MS/MS measurements of enriched N-glycoproteins from postmortem human frontal cortex tissues representing three groups:

i. individuals with no cognitive impairment and no pathological AD,
ii. asymptomatic AD subjects (pathology present without cognitive decline), and
iii. symptomatic AD subjects.

The HILIC-enriched glycopeptide runs from *PXD032219* was used. The raw LC-MS/MS files were re-analyzed using both pGlyco3 and Glyco-Decipher with the following setting to obtain glycopeptide spectral matches (GPSMs). All processed output GPSM raw files are archived in Zenodo https://zenodo.org/records/17759790 to ensure the reproducibility.

### Search engine settings

The raw files were searched using both Glyco-Decipher (version 1.0.5) and pGlyco3 (version 3.1). All searches were performed using the default parameters recommended by each search engine for intact N-glycopeptide identification. Glyco-Decipher searches employed the built-in N-glycan database containing 10,936 glycan entries, while pGlyco3 searches used the *pGlyco3-N-Human* glycan library (2,922 human N-glycans). The human proteome database *UP000005640* downloaded from UniProt on March 26, 2025. Both search engines used the same UniProtKB human reference proteome as the protein sequence database. The resulting glycopeptide spectral matches (GPSMs) from both engines were processed for subsequent analysis using glycoTraitR https://github.com/matsui-lab/glycoTraitR.

### GlycoTraitR analysis

The analysis workflow consisted of

i. importing glycopeptide spectral matches (GPSMs),
ii. extracting N-glycan structural traits and constructing the trait--GPSM matrix, and
iii. performing statistical tests to evaluate trait-level differences between clinical groups.

Below we provide representative R code that illustrates the core steps used in this case study. In addition, we compared the traits that exhibited significant changes in both the pGlyco3 and Glyco-Decipher GPSM datasets and highlight the commonly altered traits together with a brief discussion of their potential biological implications.

*<u>Import GPSM file(s)</u>*

```
library(glycoTraitR)

# Step 1: Preprocess the pGlyco3 GPSM matrix
```

```
gpsm_dir <- "/path_to_your_file/pGlycoDB-GP-FDR-Pro-Quant-
Site.txt"
pGlyco3_gpsm <- read_pGlyco3_gpsm(gpsm_dir)

# Step 1: Preprocess the GlycoDecipher GPSM folder
gpsm_folder_dir <- "/path_to_your_folder/"
decipher_gpsm <- read_decipher_gpsm(gpsm_folder_dir)

# Load sample metadata (a data frame)
meta <- readRDS("/path_to_your_file/meta.rda")
head(meta)
```

```
# A tibble: 6 × 22
  Diagnosis        `Sample number` `Tissue weight (mg)` `Age at
baseline`
  <fct>                      <dbl>                 <dbl>         <dbl>
1 Asymptomatic                  1                    62          77.1
2 Normal                      110                    72          70.8
3 Normal                      118                    74          75.7
4 Normal                       12                    57          80.8
5 Normal                      125                    52          89.2
6 Normal                      127                    24          85.2
# 18 more variables: `Age at death` <dbl>, `APOE gene` <chr>,
#   Braak <dbl>, CERAD <dbl>, `Global Cog Func` <chr>,
#   Reagan <dbl>, arteriol_scler <chr>, ci_num2_gct <dbl>,
#   ci_num2_mct <dbl>, cvda_4gp2 <chr>, dlbdx <dbl>, Sex <chr>,
#   hspath_any <dbl>, Race <chr>, tdp_cs_6reg <chr>,
#   tdp_stage4 <chr>, PMI <dbl>, file <chr>
```

*Constructing a trait-GPSM matrix*

```
# Step 2: Build trait--GPSM SummarizedExperiment object
pGlyco3_trait_se <- build_trait_se(
  pGlyco3_gpsm,
  from   = "pGlyco3",
  motifs = NULL,
  level  = "protein",
  meta   = meta
)

decipher_trait_se <- build_trait_se(
  decipher_gpsm,
  from = "decipher",
  motifs = NULL,
  level = "protein",
  meta = meta)
```

*Statistical testing to compare between clinical groups*

```
# Step 3: Perform trait-wise differential analysis
pGlyco3_res <- analyze_trait_changes(
  pGlyco3_trait_se,
  group_col    = "Diagnosis",
  group_levels = c(test = "Symptomatic", control = "Normal"),
  min_psm      = 20
)

decipher_res <- analyze_trait_changes(
  decipher_trait_se,
  group_col = "Diagnosis",
  group_levels = c(test = "Symptomatic", control = "Normal"),
  min_psm = 20)
```

*Find commonly changed trait*

```
library(magrittr)

trait_level <- merge(
  decipher_res[, 1:2],
  pGlyco3_res[, 1:2],
  by = c("trait", "level")
)

decipher_res %>%
  filter(trait %in% trait_level$trait,
         level %in% trait_level$level)
pGlyco3_res %>%
  filter(trait %in% trait_level$trait,
         level %in% trait_level$level)
```

The overlapping traits detected by GlycoDecipher were:

```
       trait    level   l_pval     f_val    t_pval     t_val
t12   Fucose     DPP6  0.05577   3.69084   0.01907   2.36091
t15 Antennas    LAMP2  0.01587   5.89225   0.02125  -2.32218
t19   Bisect    DPP10  0.06851   3.42793   0.01781   2.46415
```

The corresponding overlapping traits detected by pGlyco3 were:

```
       trait    level   l_pval     f_val    t_pval     t_val
t14   Fucose     DPP6  0.09573   2.79959   0.04507   2.02367
t16 Antennas    LAMP2  0.02312   5.21728   0.02599  -2.24018
t18   Bisect    DPP10  0.13468   2.28684   0.04432   2.06419
```

*Result*

In the case study, three trait--protein combinations were consistently highlighted by analyzing both pGlyco3 and Glyco-Decipher GPSM result: increased antennary branching on LAMP2, and reduced fucosylation and bisecting GlcNAc on DPP6 and DPP10, respectively.

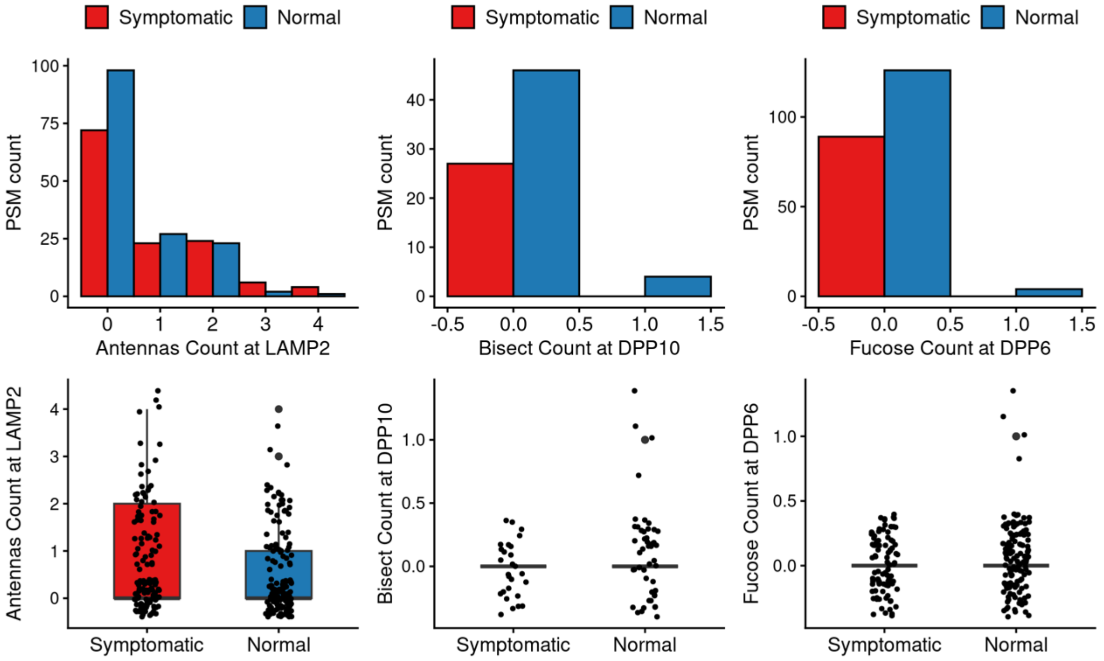


Figure S 2. Frequency histograms and boxplots of the highlighted glycan trait--protein changes for the pGlyco3 results. Symptomatic AD subjects show increased antennary branching on LAMP2 and modest reductions in bisecting GlcNAc on DPP10 and fucosylation on DPP6 compared with cognitively normal controls.

These proteins have established links to Alzheimer's disease. LAMP2, a highly glycosylated lysosomal membrane protein, is essential for autophagy and lysosomal homeostasis (Qiao et al., 2023), and its dysregulation has been reported in AD (Loeffler et al., 2018). Increased antennary complexity may reflect altered Golgi apparatus changes (Fisher et al., 2019). DPP6 and DPP10 are modulators that influence neuronal excitability. DPP6 reduction has been associated with cognitive decline (Cacace et al., 2019), while DPP10 accumulates in tau-positive pathology (Chen et al., 2014). The observed decreases in fucosylation (DPP6) and bisecting GlcNAc (DPP10) align with known AD-associated alterations in N-glycan processing pathways. These results suggest that glycan trait changes of LAMP2, DPP6, and DPP10 can be detected and may represent a new molecular layer of AD-related function. Together, these analyses highlight the utility of trait-based representations for uncovering meaningful glycosylation changes.
